# Katnip is required to maintain microtubule function and lysosomal delivery to autophagosomes and phagosomes

**DOI:** 10.1101/2022.02.22.481453

**Authors:** Georgina P. Starling, Ben A Philips, Sahana Ganesh, Jason S. King

**Affiliations:** Department of Biomedical Science, University of Sheffield

**Keywords:** Autophagy, microtubule, microtubule repair, lysosome, *Dictyostelium*, phagocytosis

## Abstract

The efficient delivery of lysosomes is essential for many cell functions, such as the degradation of unwanted intracellular components by autophagy and the killing and digestion of extracellular microbes within phagosomes. Using the amoeba *Dictyostelium discoideum* we find that cells lacking Katnip (Katanin interacting protein) have a general defect in lysosome delivery and whilst able to make autophagosomes and phagosomes correctly are then unable to digest them.

Katnip is largely unstudied yet highly conserved across evolution. Previously studies found Katnip mutations in animals cause defects in cilia structure. Here we show that Katnip plays a general role in maintaining microtubule function. We find that loss of Katnip has no overall effect on microtubule dynamics or organisation, but is important for the transport and degradation of endocytic cargos. Strikingly, Katnip mutants become highly sensitive to GFP-tubulin expression, which leads to microtubule tangles, defective anaphase extension and slow growth. Our findings establish a conserved role for Katnip in the function of all microtubules, not just cilia as previously reported. We speculate this is via a key function in microtubule repair, required to maintain endosomal trafficking and lysosomal degradation.

## Introduction

Endocytic trafficking is highly complex and dependent on a wide range of cellular machinery to generate vesicles at the correct place and time, of the correct composition, and then deliver them to where they are needed. Consequently, many elements and regulatory mechanisms remain poorly understood. Autophagy is a key endocytic pathway that allows cells to capture and digest unwanted cytosolic components. This is important for the removal of damaged or superfluous proteins and organelles and also allows cells to recycle their cytosol to regenerate energy and nutrients during starvation. Autophagy therefore plays a broad range of physiologically important roles such as preventing the accumulation of damaged components that can cause neurodegeneration and cancer and allowing cells to survive starvation (Aman et al., 2021).

Autophagosomes form by the generation of a cup-shaped structure in the cytosol, called an isolation membrane or phagophore. This expands and encircles a portion of the cytosol to generate a characteristic double-membraned vesicle. These then fuse with lysosomes leading to acidification and digestion of the captured contents. Whilst much of the basic machinery required for autophagosome generation has now been identified, our understanding of all the regulatory components and additional factors required to complete the process is incomplete.

A useful model system to investigate the autophagy pathway is the soil amoeba *Dictyostelium discoideum*. Like mammalian cells, *Dictyostelium* use autophagy to survive starvation and remove pathogens, but they also require it during their developmental cycle when ~200,000 individual cells come together and differentiate to form a multicellular fruiting body consisting of a spore head held aloft by a stalk (Mesquita et al., 2016).

The autophagy machinery in *Dictyostelium* closely represents that in mammalian cells as both form isolation membranes adjacent to the ER, in contrast to the organelle-independent phagophore assembly site (PAS) found in yeast (King et al., 2011). Consequently *Dictyostelium* possesses several genes required for autophagy in animals that have been lost in the classical genetic system *S. cerevisiae* (King, 2012; Tian et al., 2010).

To identify new components of the autophagy machinery, the Kessin group took advantage of their observation that upon starvation, autophagic degradation causes cells to deplete their cytosol and shrink over time (Otto et al., 2004). Therefore, wild-type *Dictyostelium* change buoyancy on a Percoll density gradient upon starvation, whereas autophagy deficient mutants do not. This allowed mutants unable to digest their cytosol to be isolated from a library by Percoll separation. A secondary test for defective development led to the identification of several mutants sharing both autophagy-associated phenotypes (Tekinay and Kessin, Colombia University, personal communication of unpublished screen) (Otto et al., 2003b; Otto et al., 2004). One of these was the previously unstudied gene DDB_G0275861, which we validate and characterise in this study.

DDB_G0275861 is completely unstudied, but has highly conserved orthologues throughout the eukaryotes with the notable exceptions of all fungi and the fruit fly *Drosophila melanogaster*. Orthologues can easily be recognised by the presence of a unique and highly conserved domain of unknown function - DUF4457. DUF4457 domains always occur as 3-4 tandem repeats and are not found in any other proteins (Figure 1A and Supplementary Figure 1A) (Sanders et al., 2015). The rest of the protein is more divergent, with no clear homology to any other proteins.

**Figure 1:**
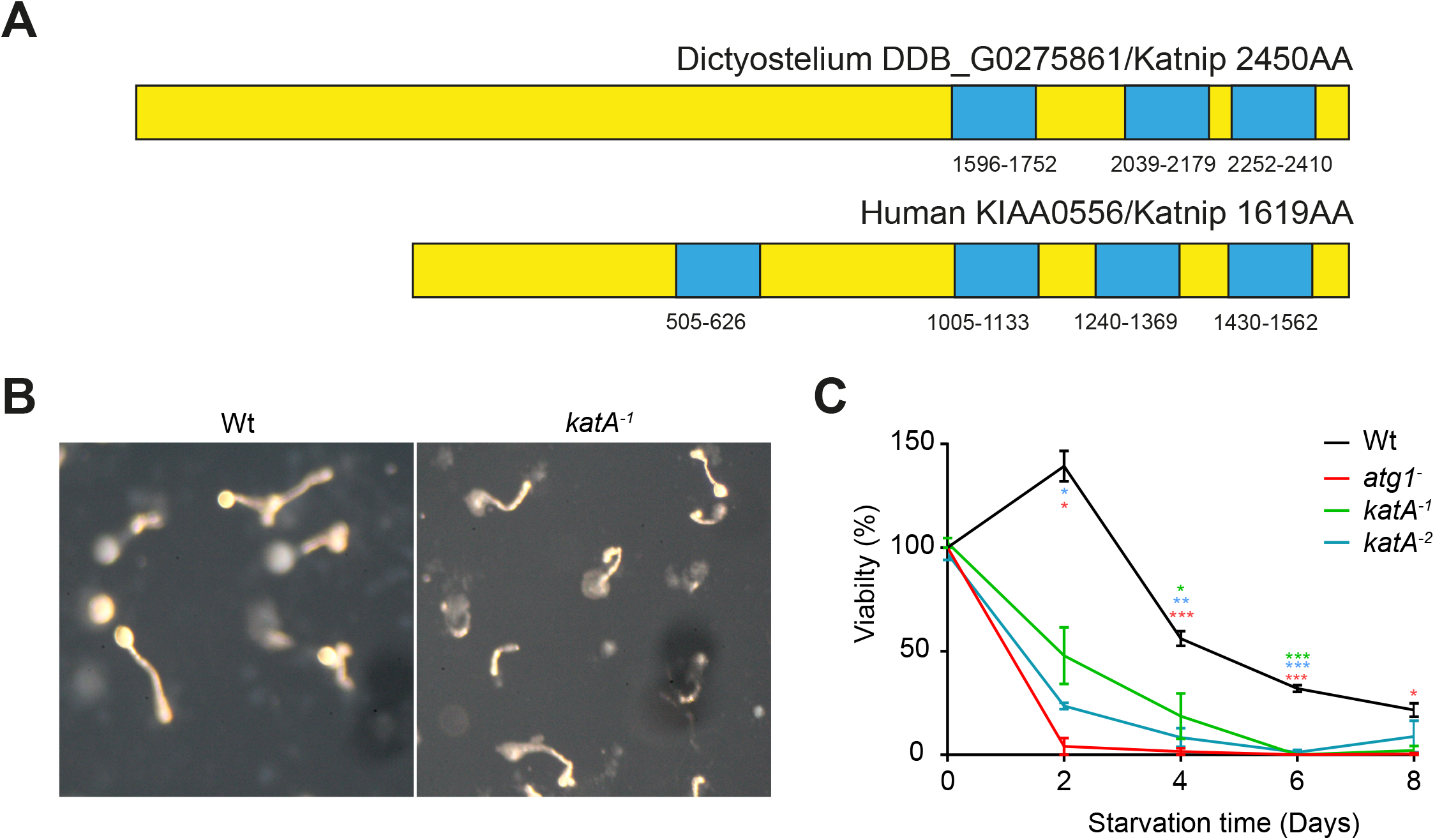
Disruption of *KatA* causes autophagy-deficient phenotypes. (A) Schematic representation of the structure of Dictyostelium and human Katnip orthologues. Blue boxes indicate the positions of the DUF4457 domains unique to this protein. (B) Development of wild-type controls and *katA*^*-1*^ mutants on nitro-cellulose filters, indicating disrupted fruiting body formation. (C) *katA*^*-*^ mutants rapidly lose viability under arginine and lysine starvation, compared to wild type controls, but less quickly than completely autophagy-deficient Atg1-null cells.

Loss of function mutations of the human orthologue (KIAA0556) were previously identified as a cause of Joubert’s syndrome, an inherited developmental disorder caused by defective cilia (Sanders et al., 2015). Cilia are microtubule-based structures that protrude from the cell surface and are involved in extracellular fluid motility, signalling and mechanosensation (Malicki and Johnson, 2017). KIAA0556 was subsequently shown to bind the microtubule severing protein Katanin, and renamed Katnip (Katanin-interacting protein) (Sanders et al., 2015). We will therefore also use Katnip to refer to the *Dictyostelium* protein, with *katA* for the corresponding gene.

Katnip was completely unstudied prior to discovery of its role in ciliopathies, and its molecular function remains unknown. In vertebrates, GFP-Katnip localises to the centrosome and cilia basal body and the primary phenotype of Katnip-deficient animals appears to be mild disruption of cilia organisation; mutant cells still form the same overall structure but have more variation in the number of microtubules within cilia, which appear slightly elongated and form at lower frequency. Consistent with a role in microtubule regulation, Katnip directly interacts with both tubulin and Katanin and high levels of overexpression cause accumulation of acetylated microtubules in retinal pigment epithelial cells (hTERT-RPE1) (Sanders et al., 2015).

How Katnip contributes to microtubule and/or cilia activity remains unknown. As *Dictyostelium* and many other organisms that possess Katnip do not produce cilia, this indicates a broader functional role that remains unexplored. Here we describe the role of Katnip (KatA) in *Dictyostelium*, indicating a new, general function in microtubule maintenance required for effective lysosomal trafficking and degradation of both autophagosomes and phagosomes.

## Results

### Katnip mutants exhibit multiple autophagy-deficient phenotypes

The original screen was performed using a library of insertional mutants generated in the DH1 genetic background (Kuspa and Loomis, 1992). We therefore started by generating new *katA* knockouts by homologous recombination to recapitulate and confirm the reported phenotypes (Supplementary Figure 1B). Multiple independent mutants were isolated in both Ax2 and Ax3 genetic backgrounds and had comparable phenotypes (Supplementary Figure 1B and C). All data presented is from the Ax2 mutant strains JSK14 and 15, referred to as *katA*^*-1*^ and *katA*^*-2*^ respectively.

We started by confirming the developmental phenotype originally used to identify *katA*-null cells as potential autophagy-defective mutants. Mutant cells and wild-type controls were therefore plated on filters and allowed to develop overnight. Whilst wildtype-cells robustly formed fruiting bodies with long stalks and large fruiting bodies, fruiting body formation was much weaker in *katA* mutants, with much shorter or collapsed stalks and almost no sorus (Figure 1B). This is not as severe as the complete loss of aggregation observed upon disruption of the single Atg1 orthologue that is essential for autophagosome formation, but is similar to that found in less penetrant autophagy mutants, such as upon disruption of one of the two Atg5 or Atg6 alleles (Otto et al., 2003b).

We then tested the ability of *katA*^−^ mutants to survive starvation, as an additional key autophagy phenotype. Recycling of nutrients by autophagic degradation of the cytosol is critical to survive starvation and autophagy deficient cells rapidly lose viability upon amino acid starvation (King et al., 2011; Otto et al., 2003b). Consistent with this, *katA*^*-*^ mutants also begun to lose viability almost immediately following arginine and lysine starvation, although not as rapidly as cells lacking Atg1 (Figure 1C). Katnip-null cells are therefore deficient in cytosolic digestion, development and surviving starvation, consistent with a functionally important role in the autophagic pathway.

### Katnip mutants have a specific defect in autophagosome degradation

To investigate where in the autophagy pathway the defect arises, we directly examined autophagosome formation using the canonical autophagy reporter GFP-Atg8a. By fluorescence microscopy Katnip-null cells still formed a comparable number of autophagosomes to wild-type controls which accumulated at a normal rate after addition of protease inhibitors to block degradation (Figure 2A and B).

**Figure 2:**
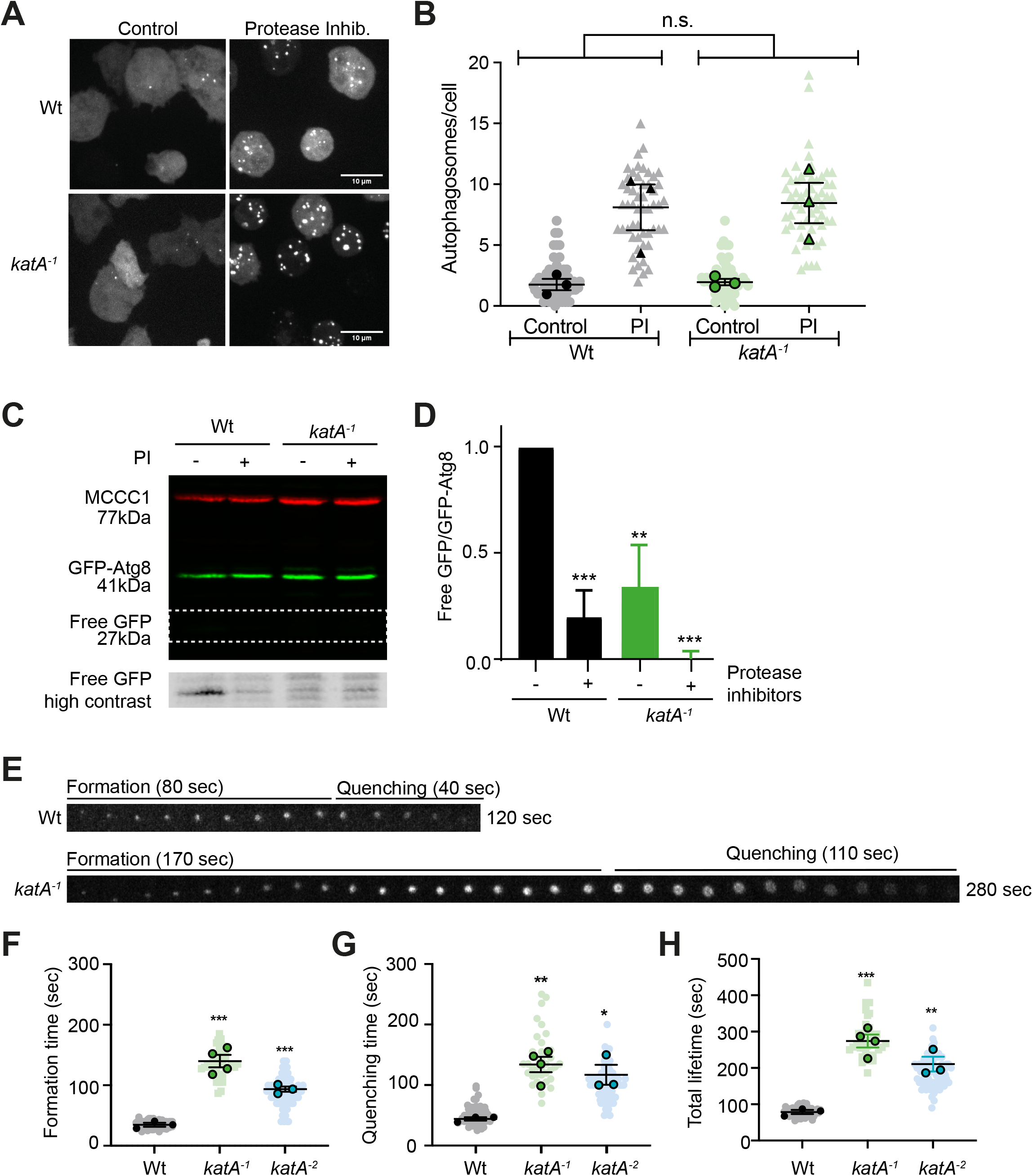
Autophagosome formation and digestion in *katA* null cells. (A) Images of cells expressing GFP-Atg8 before and after 1 hour treatment with protease inhibitors. Maximum intensity projections of confocal volumes, bar = 10 µm. The number of puncta are quantified in (B). (C) Autophagic flux assay based on the autophagic release of GFP from GFP-Atg8a. Cells were analysed with and without 1 hour protease inhibitor treatment by Western blot. The lower panel shows the boxed area after contrast enhancement to illustrate the free GFP-band. The ratio of this to total GFP-Atg8 is shown in (D) (n=3). (E) Shows representative time series of individual autophagosomes, from initiation to the quenching of GFP by acidification. See supplementary movie 1 for complete movies. Time to form a complete autophagosome, quench the GFP signal and total lifetime are quantified in (F-H) respectively. Solid-filled symbols represent biological repeats with individual cells or autophagosomes in faded colours. (***P<0.005, ** P<0.05, *P<0.01 by unpaired T-Test).

As autophagosome formation appeared normal we then tested whether there was a defect in degradation (autophagic flux). This can be determined by the accumulation of free GFP cleaved from GFP-Atg8a in autolysosomes as the compact GFP structure makes it relatively protected from lysosomal degradation (Cardenal-Munoz et al., 2017). By Western blot, a clear free GFP-band could be observed in wildtype cells that was almost completely blocked by protease inhibitors (Figure 2C and D). GFP-Atg8a cleavage was significantly decreased in *katA-*cells under all conditions, despite the cells having comparable numbers of autophagosomes (Figure 2B). We therefore conclude that loss of Katnip causes defective autophagosome proteolysis.

To follow individual autophagosomes in more detail, we observed GFP-Atg8a expressing cells compressed under agar by high-resolution timelapse microscopy. This mechanically stimulates autophagosome formation as well as reducing the Z-depth of the cells, enabling individual autophagosomes to be followed over time (King et al., 2011). This allows direct observation of phagophore expansion through to complete autophagosome formation. Subsequent lysosomal fusion and acidification is then indicated by disappearance of the GFP-signal as it becomes quenched at low pH (Supplementary Movie 1) (Mesquita et al., 2016). Whilst the autophagosomes formed by Katnip-deficient cells appeared indistinguishable from controls, both formation and acidification phases were significantly slower, taking almost three times longer to complete the cycle (Figure 2E—H). This confirms that loss of Katnip does not prevent autophagosome formation *per se*, but perturbs the underlying dynamics sufficiently to inhibit proper function and flux.

### Loss of Katnip causes a general defect in lysosomal delivery

We next tested whether the observed defects are due to a specific role for Katnip in autophagy or a general trafficking defect. *Dictyostelium* amoebae are professional phagocytes and also rely on lysosomal delivery to degrade extracellular material engulfed in phagosomes. Phagosomal proteolysis can be directly measured using reporter beads which increase in fluorescence upon hydrolysis and dequenching of conjugated DQ-Bovine Serum Albumin (DQ-BSA) (Sattler et al., 2013). This showed that disruption of Katnip also caused a 70% loss in phagosomal proteolysis compared to controls (Figure 3A). This was not due to deficient lysosomal capacity as proteolysis in cell-free lysates was unaffected and cells contained comparable levels of the acid hydrolase cathepsin D (Figure 3B-C and Supplementary Figure 1D). We therefore conclude that loss of Katnip causes a general defect in lysosomal degradation due to inefficient delivery to nascent phagosomes and autophagosomes.

**Figure 3:**
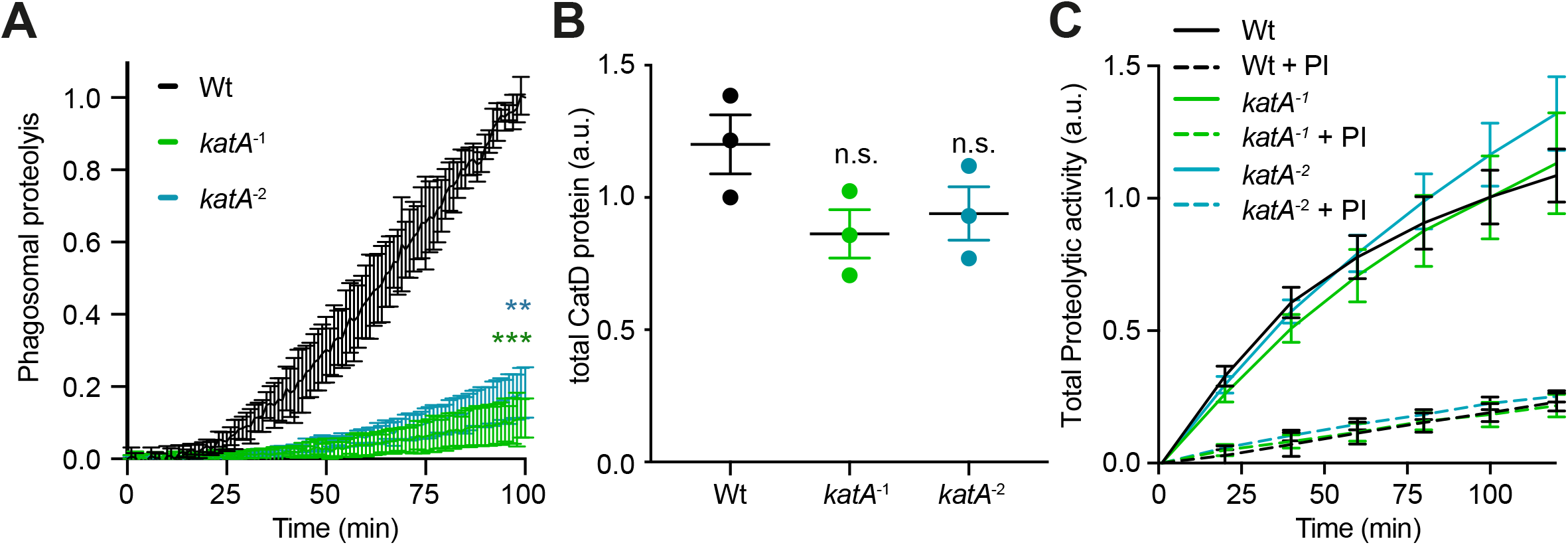
Phagosomal proteolysis is also perturbed in *katA*^*-*^ cells. (A) Proteolysis of DQ-BSA conjugated beads after phagocytosis, indicating severely inhibited digestion in *katA*^*-*^ mutants. (B) Total cathepsin D protein levels are unaffected as measured by western blot (see Supplementary Figure 1D for full blots). (C) Measurement of total proteolytic activity performed as in (A) but in cell lysates rather than intact cells. Dashed lines indicates control samples treated with protease inhibitors (PI).

### *Dictyostelium* Katnip has a conserved localisation to the centrosome and microtubules

To further understand Katnip function we examined its localisation by expression as a GFP-fusion. Unfortunately, due to the presence of a large number of repetitive sequences encoding poly-glutamine and asparagine tracts at the N-terminus of the gene we were unable to clone the full-length *Dictyostelium discoideum* gene. We therefore took advantage of the recently sequenced genome of the closely-related species *Dictyostelium lacteum* which lacks the propensity for polyN/Q repeats that so often confounds molecular biology in *D. discoideum* (Malinovska et al., 2015).

Expression of GFP-fused to *D. lacteum* Katnip (GFP-dlKatnip) in *D. discoideum* showed a significant cytosolic pool and enrichment at a single puncta near the nucleus. Additional puncta were also occasionally observed in highly expressing cells, although the perinuclear spot was always clearest. This was confirmed as the centrosome by colocalisation with the marker CP224 (Figure 4A) (Graf et al., 2000). Centrosomal localisation was maintained during mitosis, as well as an additional weaker recruitment to the spindle (Figure 4B). Serendipitously, during extended live-cell imaging we found that GFP-dlKatnip underwent a dramatic relocalisation to form large numbers of bright puncta throughout the cell with a simultaneous reduction of the cytosolic background (Figure 4C, Supplementary Movie 2). This was reproduced by treating cells with 5mM hydrogen peroxide without imaging for 30 minutes, indicating this response was due to photooxidative damage induced by imaging (Figure 4D).

**Figure 4:**
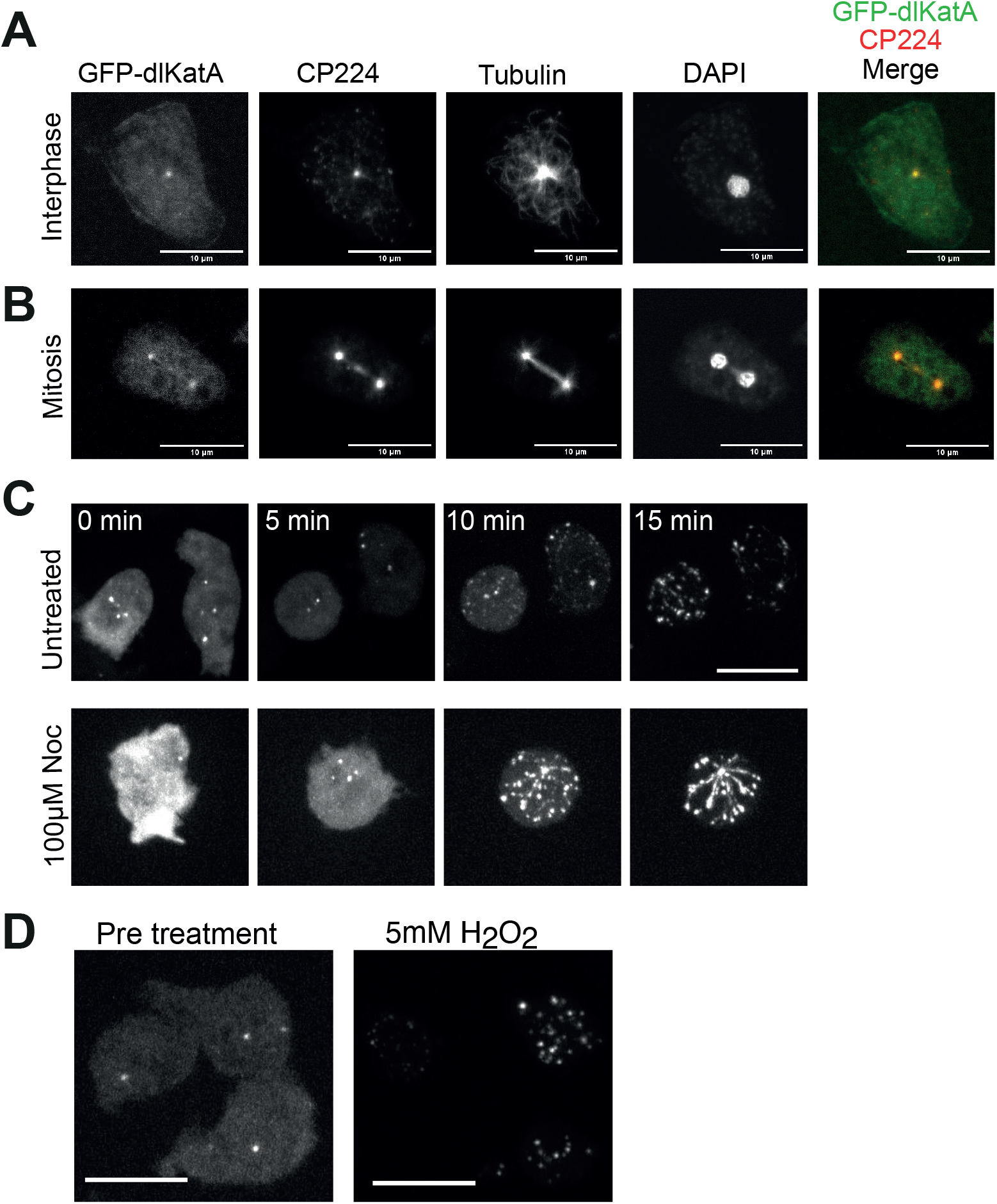
GFP-Katnip localises to centrosomes and puncta on microtubules upon oxidative damage. (A) Wild-type cells expressing GFP-dlKatA fixed and stained for the centrosomal marker CP224, tubulin and nuclei (DAPI). (B) Shows an equivalent cell during mitosis, with GFP-dlKatA localising to both the spindle poles and microtubules iin anaphase. (C) Shows frames from a timelapse of GFP-dlKatA expessing cells under prolonged illumination. GFP gradually accumulates in puncta that appear along fibrous lines. The bottom row indicates the same experiment, performed after 20 mins in 30µM nocodazole in order to reduce the microtubule array. See Supplementary Movies 2 and 3 for the full sequences (D) Shows the same cells after chemical induction of oxidati ve damage with 5mM H_2_O_2_. All scale bars indicate 10µm.

The induced GFP-dlKatnip puncta appeared to associate along filamentous structures which resembled microtubules. To confirm this, we sought to enhance this localisation by shortening the microtubule network with the microtubule depolymerising drug nocodazole prior to imaging. This shortened the microtubules and increased the density of GFP-dlKatnip puncta labelling, clearly highlighting the microtubule network (Figure 4C, bottom panels and Supplementary Movie 3). Whilst the physiological significance and mechanism driving this relocalisation is unclear, a similar localisation of GFP-Katnip puncta along microtubules was described in mammalian cells upon high levels of overexpression, and Katnip directly binds to tubulin (Sanders et al., 2015). This implies conserved function and suggests a potential conditional ability to bind along microtubules that becomes constitutive upon extreme overexpression as a GFP fusion and/or oxidative damage.

### Katnip is required to tolerate GFP-tubulin expression

We next investigated how loss of Katnip affected the microtubule cytoskeleton by expression of GFP-α-tubulin. *Dictyostelium* microtubules are highly stable and form exclusively from the centrosome to generate a single array that is evenly distributed throughout the cytosol.

In *katA*-null cells GFP-tubulin revealed a general disruption of the microtubule organisation, with 30% of cells exhibiting highly stable tangles and/or a collapsed array that left the majority of the cytosol microtubule-free (Figure 5A and B and Supplementary Movie 4). Surprisingly however, this phenotype appeared to be specifically induced by GFP-fused α-tubulin as *katA-*cells stained for endogenous tubulin had no significant increase in aberrant microtubule arrays (Figure 5C).

**Figure 5:**
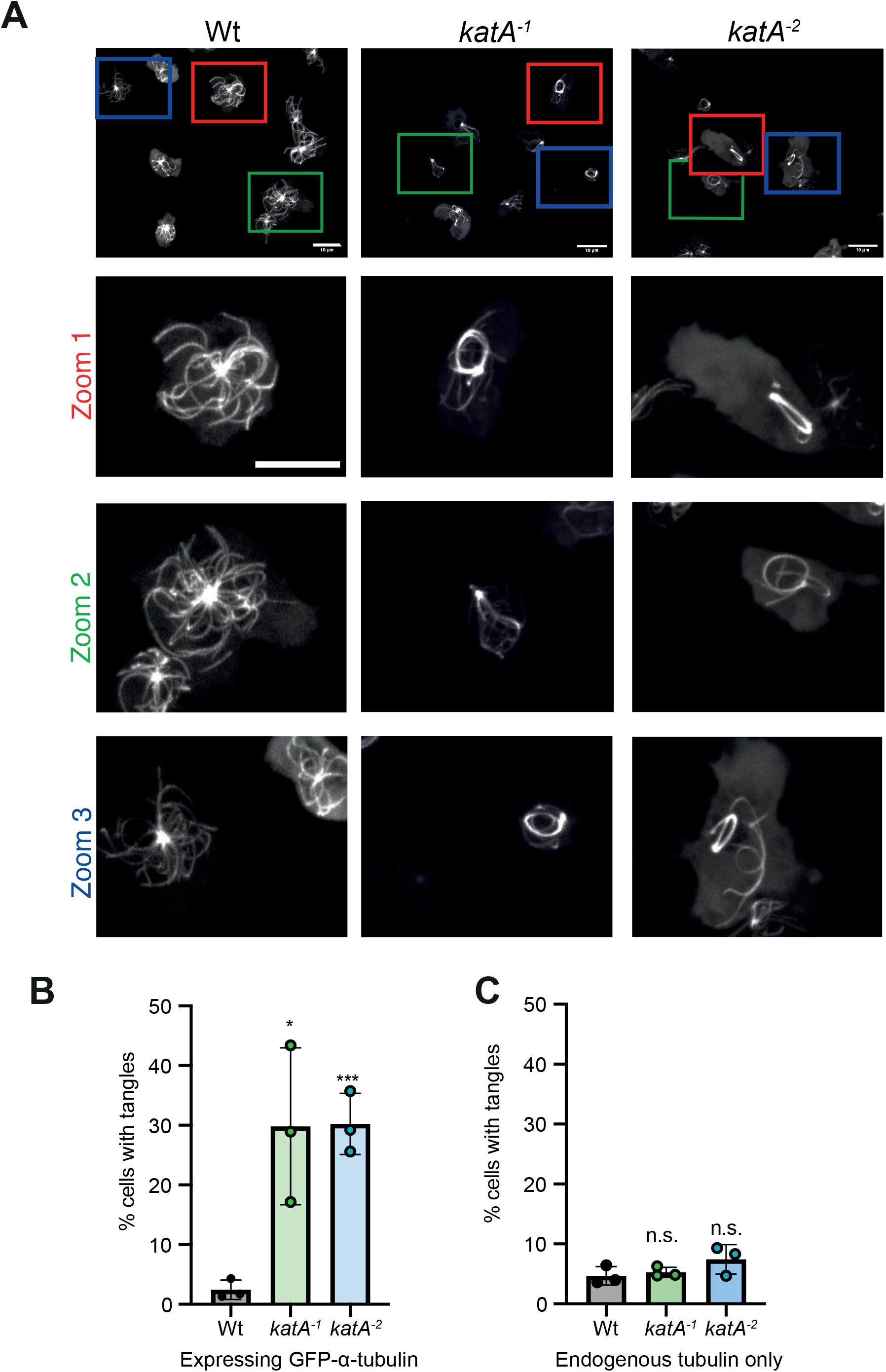
GFP-α-tubulin causes the formation of microtubule tangles in *katA* null cells. (A) Shows maximum intensity projections from confocal sections of cells expressing GFP-α-tubulin. Three sections from each field of view are enlarged below as indicated by their coloured boxes (Scale bars =10µm). The proportion of cells line with tangles upon GFP-α-tubulin expression from 3 independent experiments is shown in (B). (C) Shows the equivalent analysis of untransformed cells, fixed and stained for endogenous tubulin, indicating the tangle phenotype is dependent on GFP-α-tubulin expression. Timelapses of these tangles are shown in Supplementary M

GFP-α-tubulin expression is used widely and had no significant effect on the growth or microtubule organisation of wild-type *Dictyostelium* (Figure 6A). Untransformed *katA-*cells also grew at a normal rate, but GFP-α-tubulin expression caused their generation time to increase from 15 to 48 hours (Figure 6A, see Supplementary Figure 2A for complete growth curves). This defect was not due to a defect in cytokinesis as all the cell lines had comparable levels of multinucleate cells (Supplementary Figure 2B).

**Figure 6:**
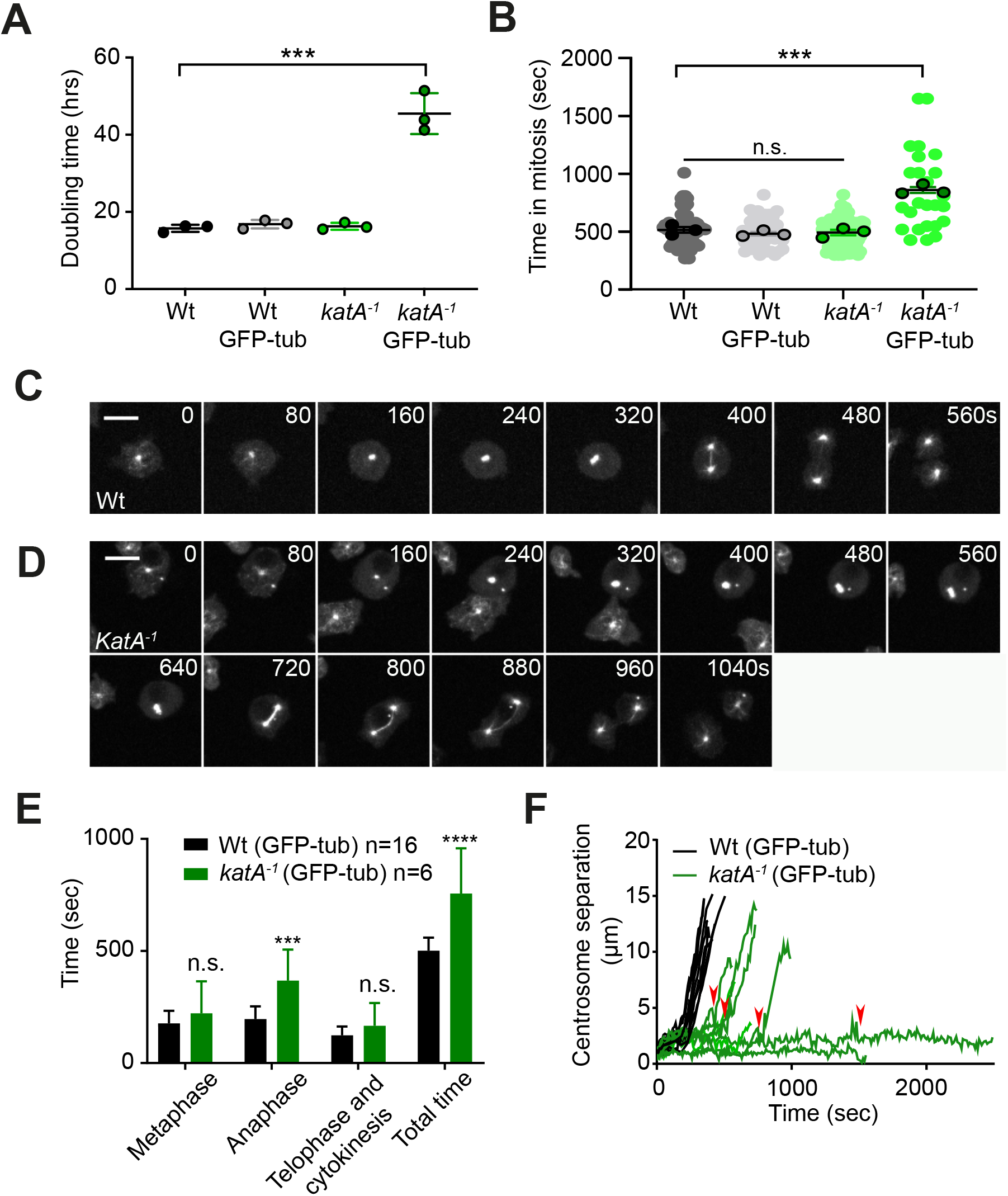
GFP-α-tubulin specifically disrupts both growth and anaphase in *katA*^*-*^ cells. (A) Growth rate of cells with and without GFP-α-tubulin expression in standard HL5 axenic medium. (B) Time spent in mitosis, estimated by cell rounding during bright-field timelapse microscopy. (C) and (D) show confocal time series of GFP-α-tubulin dynamics during mitosis of wild-type and *katA*^*-1*^ cells respectively. The full time series is shown in Supplementary Movies 5 and 6. The time spent in each phase of mitosis is shown in (E). (F) Shows the length of the mitotic spindle over time from timelapse movies. Each line represents an individual cell division. Points where spindle elongation stalls or retracts are indicated by red arrows. All scale bars =10µm, *** P<0.005 (un-paired T-Test.

The 3-fold decrease in growth rate induced by GFP-α-tubulin cannot be solely due to the 30% of *katA-* cells that contain tangles. Tangles are therefore likely to be extreme manifestations of a general GFP-α-tubulin intolerance across the population. Despite the presence of the bulky fluorescent tag, GFP-α-tubulin is generally well-tolerated in most organisms. However high expression levels have been reported to induce mild cold-intolerance in *S. cerevisiae* (Carminati and Stearns, 1997) and a decrease in microtubule number in *Dictyostelium* (Kimble et al., 2000). The increased sensitivity of *Dictyostelium* cells lacking Katnip to GFP-α-tubulin indicates either a general role for Katnip in microtubule regulation or a more specific role in tolerating microtubule stress.

### GFP-tubulin causes a specific defect in spindle elongation in Katnip-null cells

To better understand how GFP-tubulin expression affects microtubule dynamics and growth of Katnip mutants we examined mitosis as a classical microtubule-driven process. Using brightfield imaging to observe division with minimal phototoxicity indicated that whilst most cells still successfully divided, expression of GFP-α-tubulin specifically increased the variability and average time spent in mitosis in cells lacking Katnip (Figure 6B). More detailed analysis, observing GFP-α-tubulin dynamics by confocal microscopy indicated a specific defect during anaphase, when the microtubule spindles pull the centrosomes apart (Figure 6C-E, Supplementary Movies 5 and 6). Whilst wild-type spindles extended in a steady and highly reproducible manner, *katA*-null spindles frequently stalled mid-elongation and either paused or retracted (Figure 6F). This resulted in a significantly longer and more variable time in anaphase and an overall doubling of the average time in mitosis, confirming our observations with brightfield microscopy.

These experiments demonstrate that Katnip is required for cells to tolerate GFP-α-tubulin. Whilst this is particularly apparent during anaphase when microtubules are most mechanically active, the short proportion of time spent in mitosis means this will only make a minor contribution to the overall 3-fold increase in generation time. Katnip must therefore also play a general role in maintaining microtubule activity during interphase.

### Katnip does not regulate microtubule stability or dynamics

Microtubules are highly dynamic and their rates of polymerisation and depolymerisation are carefully controlled by a host of regulatory factors. *Dictyostelium* microtubules are relatively stable and rarely undergo the classical catastrophe/repolymerisation cycles prominent in their mammalian counterparts (Gräf, 2015). However, they still undergo dynamic instability and rapidly depolymerise in the presence of free tubulin sequestering drugs such as nocodazole. We therefore measured sensitivity to nocodazole to test whether loss of Katnip affects the underlying microtubule polymerisation dynamics.

Nocodazole sensitivity was tested by measuring changes in the maximal extent of the microtubule array over time. Using GFP-α-tubulin expressing cells, the proportion of wild-type cells covered by the microtubule array reduced from 67 ± 2.3% to 35 ±12% within 10 minutes of 30 µM nocodazole addition (Figure 7A). The effects of nocodazole are reversible, and the size of the microtubule array was fully restored within 10 minutes of drug removal (Figure 7B). Loss of Katnip had no effect on either the depolymerisation or regrowth of microtubules in this assay. To ensure any defects were not obscured due to excessive drug concentrations, the lack of effect was confirmed under conditions of partial depolymerisation using 10 µM nocodazole (Figure 7C).

**Figure 7:**
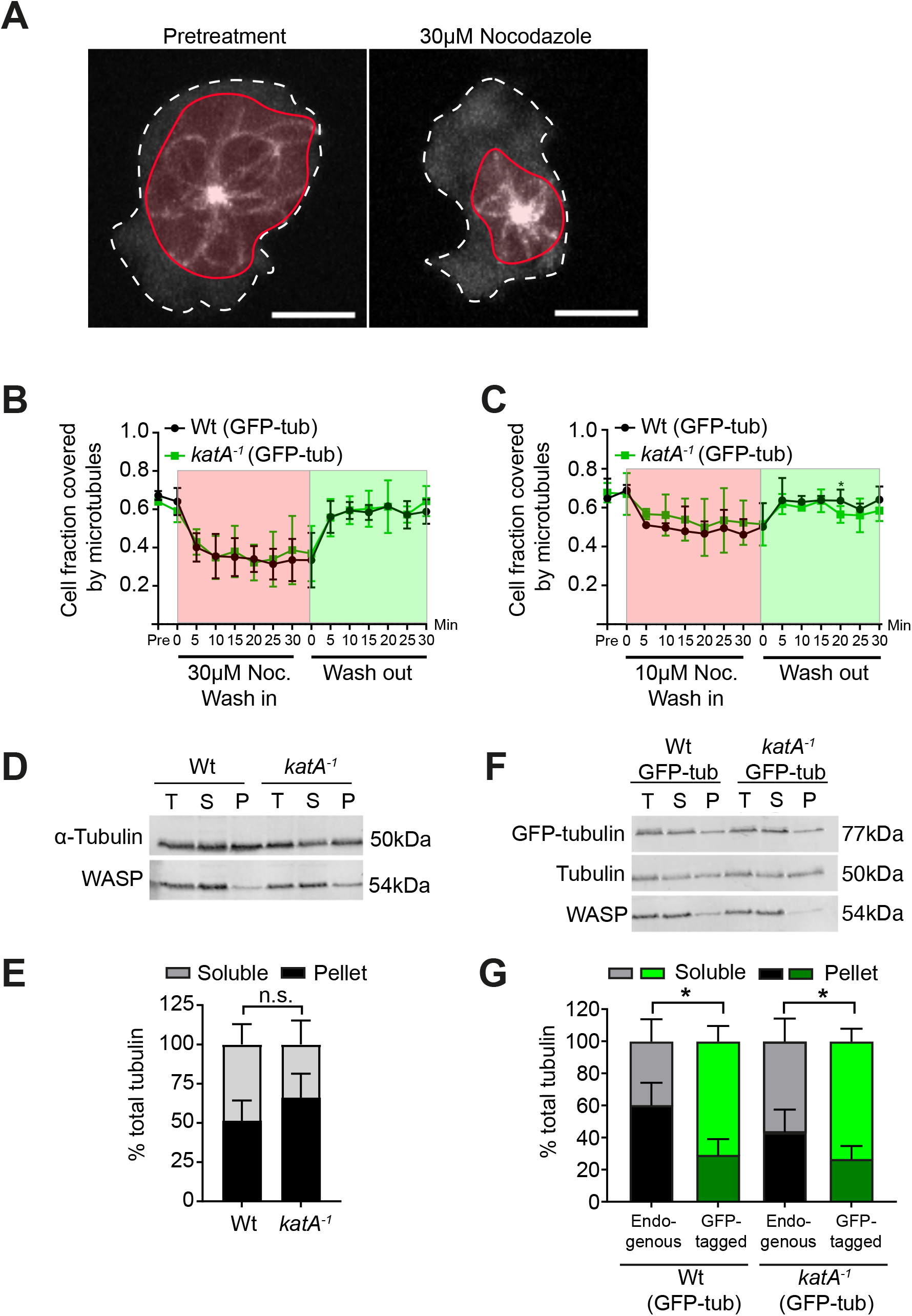
Katnip does not affect general microtubule stability. (A) Schematic for calculating the proportion of the cell area occupied by the microtubule array. Both the cell perimeter and extent of the microtubules were manually drawn from maximum intensity projections of cells expressing GFP-α-tubulin (scale bar = 5µm) (B) Shows the change in proportion of the cell covered by the microtubule array after addition and washout of 30µM nocodazole. (C) Shows the same experiment repeated with 10µM nocodazole to induce partial microtubule depolymerisation. No significant differences between wild-type and *katA*^*-*^ cells were found at any point. (D) Shows separation of the soluble (free) and insoluble polymerised) tubulin fractions from untransformed cells, analysed by western blot. WASP was used as a soluble protein control. Quantification from 3 independent experiments is shown in (E). (F) and (G) are the same experiment performed in cells expressing GFP-α-tubulin. Both endogenous and GFP-tagged α-tubulin are quantified in (G) showing GFP-α-tubulin is less efficiently incorporated into microtubules (* P<0.01, unpaired T-test). No significant differences were found between wild-type and *katA*^*-*^ cells.

As a parallel approach, we tested whether loss of Katnip affected the proportion of polymerised tubulin by fractionating polymerised (insoluble) and free (soluble) tubulin by centrifugation and subsequent Western blotting (Kimble et al., 2000) (Figure 7D-G). The predominantly soluble Wiskott Aldrich Syndrome protein (WASP) was used as a fractionation control. In this assay, loss of KatA had no significant impact on the levels of either total or polymerised endogenous α-tubulin (Figure 7D and E, cells not expressing GFP-α-tubulin). Upon transformation, both cell lines expressed GFP-α-tubulin at comparable levels (Supplementary Figure 2C) and consistent with previous reports, GFP-α-tubulin was less efficiently incorporated into microtubules than endogenous tubulin (Figure 7F and G) (Kimble et al., 2000). However, no significant changes in the polymerised fraction of either endogenous or GFP-tagged tubulin were observed in *katA*^−^ cells (Figure 7D-G). Katnip therefore does not appear to be a general regulator of microtubule assembly or disassembly and must maintain lysosomal function and tolerance of GFP-α-tubulin through an alternative mechanism.

### Katnip is required for efficient microtubule-based trafficking

Because the overall structure and stability of the microtubule array was unperturbed by loss of Katnip in the absence of exogenous GFP-α-tubulin, we next investigated whether microtubule function also remained intact. The primary role of microtubules is to act as tracks for molecular motors such as dyneins and kinesins to transport cargo around the cell. This is particularly important for endocytic trafficking and is required for lysosomal delivery to both autophagosomes and phagosomes.

The ability of phagosomes to move along microtubules can be directly observed by timelapse microscopy of 1µm latex beads after engulfment and compression of the cells to prevent migration (Rai et al., 2016). During early maturation, phagosomes move in a bi-directional manner with runs of fast movement (Rai et al., 2016). We therefore performed timelapse imaging of cells from 10 minutes after addition of beads and measured the average speed of phagosome movement (Figure 8A and Supplementary Movie 7). This showed that average speed was significantly decreased in *katA*^*-*^ cells from 0.24 ± 0.015 µm/s in controls to 0.12 ± 0.04 µm/s cells (p<0.01, T-test). To visualise these changes over time, the data were also plotted as a heat map (Figure 8B). This showed that whilst phagosomes in katA^−^ cells were able to reach similar peak speeds as wild-type controls (~1.5 µm/s), this was rarely maintained for periods longer than a few seconds, and much of the time the phagosomes remained virtually immobile.

**Figure 8:**
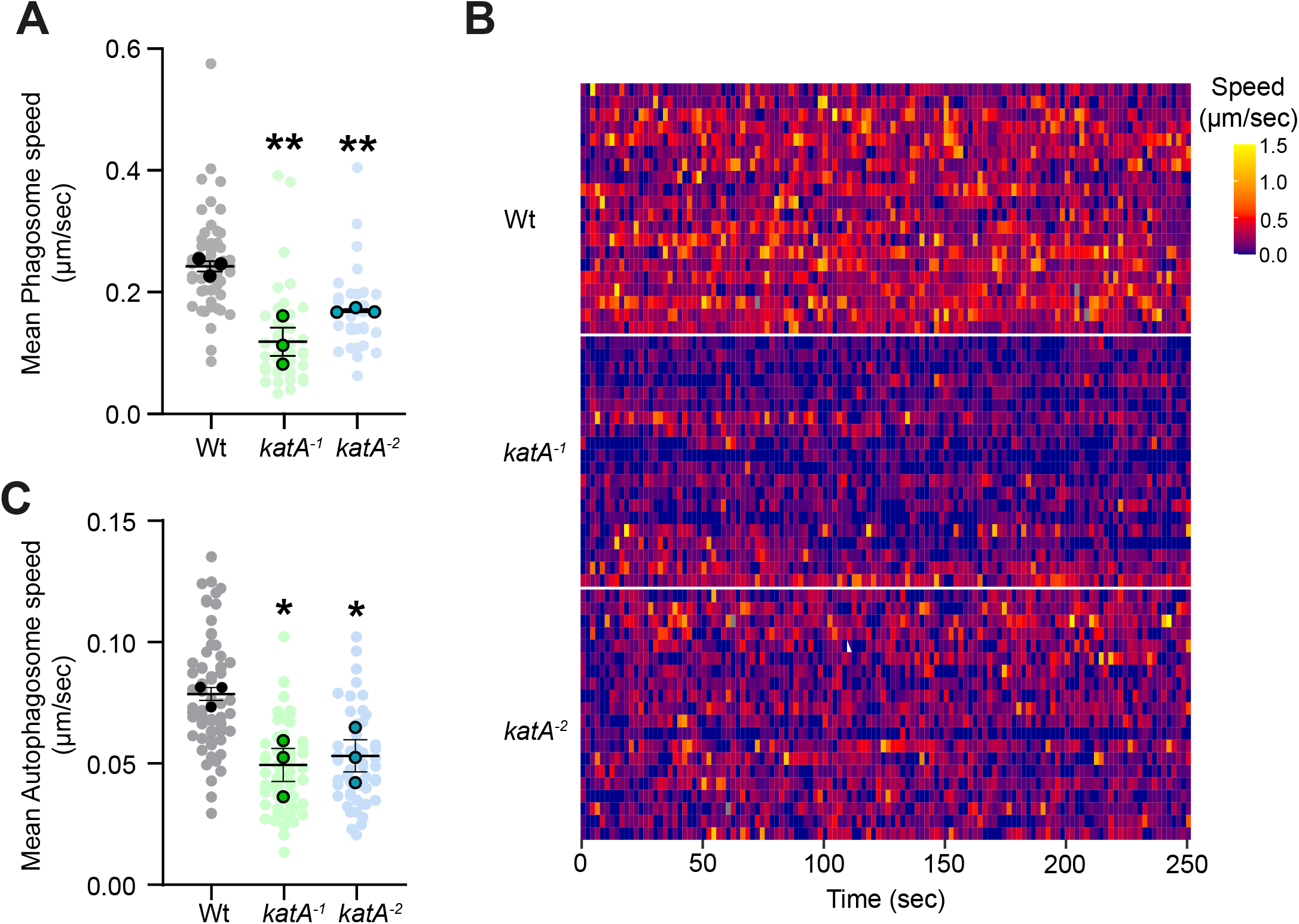
Katnip is important for efficient phagosome and autophagosome transport. (A) Shows quantification of the average speed of phagosomes moving within the cell, measured from timelapse movies after engulfment (see Supplementary movie 7 for an example). (B) Shows the same data over time plotted as a heat map. Each row represents an individual phagosome over time with colour representing its instantaneous speed. Yellow areas indicate runs of fast movement with darker blue patches indicating stalls or stoppages. A representative 20 phagosomes are shown for each cell line. (C) Shows analysis of autophagosome movement during formation and quenching, quantified from movies as in Figure 2 and Supplementary movie 1. (**P<0.05, * P<0.01, unpaired T-Test).

To establish whether autophagosome movement was also affected, we tracking their movement from the movies of compressed GFP-Atg8a expressing cells shown in Figure 2 and Supplementary Movie 1. This showed that autophagosomes also had a comparable decrease in motility in Katnip-deficient cells (Figure 8C). Therefore, absence of Katnip causes a general defect in endocytic transport, even without GFP-α-tubulin. Although how Katnip maintains microtubule-based transport remain unclear, this provides a clear mechanism for the defective lysosomal degradation of both autophagosomes and phagosomes observed in Katnip-null cells. We therefore describe a new, universal role for Katnip in microtubule function and degradation.

## Discussion

Although initially identified in a screen for autophagy-related genes, the experiments above indicate that Katnip plays a general role in ensuring lysosomal delivery to multiple compartments. Both our data and that in previous studies implicate Katnip as a key regulator of microtubule function. Although the exact role of Katnip remains unclear, an inability to use microtubules to transport vesicles to where they are needed offers a clear mechanism for the observed defects in digestion.

What role might Katnip play in microtubule transport? Microtubules and the motors that transport cargo along them are regulated in many ways. A defect in trafficking, in the absence of any major changes in microtubule organisation (in the absence of GFP-tubulin) and rates of assembly and disassembly indicates Katnip may be more important in maintenance, rather than general microtubule dynamics. Katnip localisation also hints at a general role in microtubule maintenance: whilst low levels of GFP-Katnip expression primarily localise to the centrosome in both *Dictyostelium* and mammalian cells, it can also bind along the length of the microtubule upon either overexpression or oxidative damage (Sanders et al., 2015). In highly expressing RPE1 (Retinal pigment epithelial) cells, GFP-Katnip microtubule localisation also led to increased microtubule stabilisation and acetylation (Sanders et al., 2015). Whilst the physiological relevance of overexpressed GFP-fusions or oxidative relocation is unclear, this does indicate a conditional capacity to bind the tubule body.

The clearest evidence for a role in maintaining microtubule integrity is the striking increased sensitivity to GFP-α-tubulin expression observed when Katnip is disrupted. Microtubules are formed of a highly-organised lattice of α/β tubulin dimers. It is perhaps surprising that attachment of a large GFP-moiety to these subunits is generally tolerated so well as to our knowledge no deleterious effects of GFP-α-tubulin overexpression have been reported in mammalian cells. No functional defects have previously been described in *Dictyostelium* either, despite the less efficient integration of GFP-α-tubulin into microtubules and reduced microtubule number (Kimble et al., 2000). In contrast, GFP-α-tubulin overexpression has reported to be deleterious in both *S. cerevisiae* and *S. pombe* (Carminati and Stearns, 1997; Ding et al., 1998). Like all fungi, both lack Katnip orthologues due to secondary loss through evolution.

Our observations are consistent with the hypothesis that GFP-α-tubulin expression disrupts microtubule structure, but in a way that is normally tolerated in a Katnip-dependent manner. Recent studies have highlighted that the microtubule lattice often becomes damaged when exposed to physical force, including motor walking, bending and shear forces (Schaedel et al., 2015; Triclin et al., 2021). This manifests as defects in the microtubule lattice such as the formation of holes and changes in protofilament number (Chretien et al., 1992).

Importantly, it has also been proposed that defects in the lattice structure will cause motors to stall or fall off, inhibiting endocytic transport (Liang et al., 2016). However, whilst there is now clear evidence that microtubules can be repaired through insertion of new GTP-tubulin molecules, how this is regulated remains poorly understood (Aumeier et al., 2016; Dimitrov et al., 2008; Triclin et al., 2021). One proposed mechanism is to sever damaged filaments using the activity of Katanin (Diaz-Valencia et al., 2011), a direct binding partner of Katnip (Sanders et al., 2015).

Cilia represent some of the most mechanically-exposed and long lived microtubules in animals (Schaedel et al., 2015). This is consistent with why structural defects in these are the primary phenotype in Katnip-deficient patients and animal models (Sanders et al., 2015). Microtubules are also under their greatest strain during anaphase in mitosis, where they need to pull the chromatids apart. This is again where we observe the most significant defect in *Dictyostelium*. The unusually high microtubule stability and lack of catastrophes in *Dictyostelium* may also contribute to damage accumulation over time and cause these cells to be particularly dependent on Katnip for repair.

Studies of microtubule repair are in their infancy and current lack of tools to observe microtubule ultrastructure and damage in cells makes testing these hypotheses challenging. Nonetheless, the data presented here demonstrate a conserved role for Katnip in regulating microtubule function beyond just cilia. We show this is important to maintain microtubule-based endocytic trafficking and in particular the digestion of autophagosomes and phagosomes.

Whilst there remain very few studies of Katnip function, it has a growing association with human disease. In addition to several studies finding Katnip mutations in patients with Joubert’s syndrome (Cauley et al., 2019; Niceta et al., 2020; Sanders et al., 2015), reduced expression was also found to be associated with Alzheimer’s disease (Andres-Benito et al., 2018). Heterozygous mutations in Katnip were also recently associated with hypothalamic hamartoma tumours (Fujita et al., 2019). Whether these disease associations are due to defects in cilia or endocytic trafficking, such as perturbed autophagy remains to be determined, but this work demonstrates that Katnip is fundamentally important for microtubule function, and general defects in trafficking and lysosomal degradation will have wide-ranging effects on the cell.

## Materials and Methods

### Cell culture

All experiments were performed using the *Dictyostelium discoideum* cells from the Ax2 axenic mutant background (strain ID DBS0235521). For general culture, cells were grown in sterile HL5 media (Formedium, UK) at 22°C. Starvation was performed using SIH defined medium containing or lacking Arg/Lys (Formedium) as previously described (King et al., 2011). Ax2 background *Atg1* mutants were previously described (DBS0350450) (King et al., 2013). Cells were transformed using standard *Dictyostelium* electroporation methods and transformants selected 24 hours after electroporation with either 10µg/mL, G418 10µg/mL blastidicin or 20ug/mL hygromycin.

To observe development, 10^7^ cells were washed twice before resuspension in 1mL KK2 starvation buffer (0.1M potassium phosphate pH6.1). This was then spread onto 47mm nitrocellulose filters (Millipore) on top of 3 filter paper discs pre-soaked in KK2. Excess liquid was then removed by pipetting from the bottom of the filter papers and cells were left for 24 hours before imaging.

Viability under starvation was performed based upon (Otto et al., 2003a). Cells were washed twice, resuspended at 5 × 10^5^/mL in SIH - Arg/Lys and maintained in shaking flasks. At each timepoint, volumes equivalent to 100 starting cells were spread on lawns of *Klebsiella aerogenes* on duplicate SM agar plates (Formedium). The number of colonies formed was then counted after 4-5 days.

### Plasmid constructs and transformation

The KatA knockout construct was generated by amplifying and cloning a 5.5Kb fragment of *Dictyostelium* KatA using the primers AGCAGAAGAAATAGCAGTAGTGGC / TTATTTTAAATAACTAGCCATTTCTGTAT and inserting the loxP-blasticidin resistance cassette from pLPBLP (Faix et al., 2004) as a BamHI/HindIII fragment into the same sites, deleting a 2.5Kb of coding sequence. The KO cassette was then excised and used to transform Ax2 cells. Correct insertion of the BSR cassette was screened by PCR, using the following primers, as indicated in Supplementary Figure 1B. P1=GAATCAATAACCCCAAGAGAGTTG; P2=TTCGGCAGTACATATTGAAGCG; P3=CGCTTCAATATGTACTGCCGAA; P4=CAGTTGAAGCTCCTTTCATTCC.

Full length *katA* from *Dictyostelium lacteum* was amplified from genomic DNA extracted from cells (kindly provided by Pauline Schaap) using primers tgatcaATGTATGCACTTTTTAAATTATGTTTC / tctagaTTATTTTTGGAATAATATTGATTGGGAG, also adding 5’BclI and 3’XbaI restriction sites. This was subcloned into the BglII/SpeI sites of *D. discoideum* expression vector pDM448, adding an N-terminal GFP tag (Veltman et al., 2009). GFP-tubulin and GFP-Atg8aexpression vectors (pJSK336 and pDM430 respectively) were generated previously (King et al., 2010; King et al., 2011).

### Proteolysis assays

Phagosome proteolysis was determined based on the dequenching of DQ-BSA/Alexa594 conjugated 3μm silica beads (Kisker Biotech) as described (Sattler et al., 2013). Cells were seeded at 3 × 10^5^ per well of a glass-bottom 96 well-plate (Greiner) in Lo-Flo medium (Formedium) in triplicate. After 1 hour, 1.5 × 10^4^ reporter beads were added to each well except controls and plates were spun at 800 x *g* for 10 seconds to synchronize bead uptake. Cells were then washed twice with Lo-Flo to remove unengulfed beads before fluorescence measurements at 480/510nm and 560/620nm using a plate reader every 2 minutes. After subtraction of background fluorescence, DQ-BSA Signal was normalised for bead uptake using the Alexa594 signal.

Total proteolytic activity was performed in a similar way by adding the same reporter beads to cell lysates (Buckley et al., 2019). 4 × 10^7^ cells/mL in 150mM potassium acetate pH4.0 were lysed by two freeze-thaw cycles in liquid nitrogen. Cell debris was removed by centrifugation at 18,000 x *g* for 5 minutes at 4°C before 1.23 × 10^5^ beads were added 100µL of supernatant in 96 well plates and measured as above. Cathepsin D levels were measured by Western blot using the previously published antibody (Journet et al., 1999).

### Autophagy assays

Timelapse analysis of autophagosome formation was performed using cells compressed under agar, to both improve imaging and induce autophagy via mechanical stress (Dominguez-Martin et al., 2017). Cells expressing GFP-Atg8a(pDM430) were seeded in 35mm glass bottom dishes (MatTek) and left for 1 hour to adhere. Gels were prepared by casting 10mL of 1% agarose in HL5 medium in a 10cm petri dish before excising a ~1cm^2^ slice, which was placed upon the cells and the media removed. Residual media was removed by filter paper touching the agarose square. Cells were then imaged on Perkin-Elmer Ultraview VOX spinning disk confocal microscope with x60/1.4NA objective, taking 5 optical sections at 1μm Z-spacing every 10 seconds. Autophagosomes were tracked manually from maximum intensity projections. Autophagosome closure was defined as the brightest timepoint for each autophagosome.

Autophagic flux was measured based on the lysosomal cleavage of GFP-Atg8a as described in (Cardenal-Munoz et al., 2017). 4 × 10^6^ cells expressing GFP-Atg8a were seeded in duplicate in 3mL of HL5 in 6 well plates and placed on an orbital shaker at 180rpm at 22°C. After 2 hours, cOmplete protease inhibitor cocktail (Roche) was added to one set of wells at 4x concentration. After a further hour 50µL cell suspension was transferred to a cloning ring on a 35mm glass bottomed dish (MatTek) for microscopy, and imaged as above taking single z-stacks at 0.5 µm Z-spacing. The rest of the culture was pelleted and flash frozen in liquid nitrogen before analysis by western blot. Fluorescently-labelled streptavidin, which binds only 3-methylcrotonyl-CoA carboxylase (MCCC1), the only substantially biotinylated endogenous protein was used as a loading control (Davidson et al., 2013).

### Photodamaging cells

To induce photodamage, cells expressing GFP-dlKatA were imaged continuously on a Perkin-Elmer Ultraview VOX spinning disk confocal microscope with a x100 1.4NA oil immersion objective using 488 nm laser excitation at 50% power.

### Microtubule depolymerisation assay

Cells expressing GFP-α-tubulin were seeded microscopy dishes and left 1 hour to settle. Media was then removed from the dish and replaced with either 10µM or 30µM nocodazole (Acros Organic) in HL5. New fields of view were then imaged every 5 minutes for 30 minutes before the media was again removed, and cells washed twice and replaced with HL5 without nocodazole. Cells were again imaged every 5 minutes for a further 30 minutes. Full-depth Z-stacks (0.5µm spacing) were obtained on a spinning disk microscope before quantification from maximum intensity projections using ImageJ. The area covered by microtubules was manually measured by drawing a polygon around the ends of the array and calculated as a proportion of the whole cell area. This was calculated for all complete cells in the field of view for each time point with the means proportion from each time point averaged across 3 biological repeats.

### Microtubule fractionation

Protocol was adapted from (Kimble et al., 2000). 2 × 10^7^ cells were washed once in KK2 (0.1M potassium phosphate pH 6.1, before pelleting (50 x *g* for 2 minutes) and resuspension in 200µL HEMS buffer (50mM HEPES (Sigma Aldrich), 2mM EGTA, 5mM Mg-acetate (Fisher), 10% Sucrose (Fisher), 2% Triton X-100 (Fisher), at pH7.4) with 1% HALT protease inhibitors (Thermo Fisher). After 2 minutes at room temperature, 30µL of lysate was removed for the ‘total tubulin’ fraction before the insoluble polymerised and free tubulin fractions were separated by centrifugation at 2700 x *g* for 1 minute. Samples were then analysed by western blot, using antibodies against β-tubulin (YL1/2, gift from Ralph Graf), GFP (rabbit polyclonal, gift from Andrew Peden), or WASP (loading control, gift from Robert Insall).

### Mitosis imaging

Cells were seeded in HL5 at 0.5 × 10^5^ in 35mm glass dishes and left overnight before imaging. Brightfield movies were obtained taking images every 30 seconds for 2 hours on a LD A-plan x20 air objective on Zeiss Axiovert widefield microscope. Mitosis was timed from initial rounding until daughter cell scission. To image microtubules, cells expressing GFP-tubulin were imaged under brightfield and 488 laser every 10 seconds for 10 to 40 minutes on Perkin-Elmer Spinning Disk Confocal microscope with an 60x/1.4NA oil immersion objective. Total mitosis time of each cell was plotted for each cell line and a T-test performed between cell lines. Distance between centrosomes was measured during anaphase manually using ImageJ.

### Immunofluorescence

*Dictyostelium* cells seeded on acid-washed coverslips at 2 × 10^5^/mL were fixed in ultracold methanol as described in (Hagedorn et al., 2006). After blocking with 3% BSA in PBS for 30 minutes, coverslips were incubated with primary antibody (YL1/2 for tubulin staining, CP224 from the Graf lab) diluted in 3%BSA in PBS for 30 minutes and then repeated with secondary antibody (Anti Rat 448/594, Anti Rabbit 594 or Anti Mouse 594 all from Life Sciences). Coverslips were washed and then mounted with Prolong Gold (Invitrogen) with DAPI. Slides were left overnight to harden and imaged by either spinning disk or Zeiss Airyscan confocal microscopes using 60x objectives.

### Vesicle motility assays

Phagosome motility was measured using the assay described by (Rai et al., 2016). 2 × 10^6^ cells were seeded in HL5 medium in glass-bottomed microscopy dishes. 2 × 10^7^ 1µm YG-labelled polystyrene beads (Polysciences 15702-10) were then added, swirled to mix and left for 10 minute to be engulfed. Cells were then overlaid with a thin sheet of 2% agarose in HL5 medium as per imaging autophagosomes above. Movies were then taken using Differential interference contrast (DIC) microscopy, capturing 200 frames at 2 second intervals.

Movement of both autophagosomes and phagosomes was tracked manually using the mTrackJ plugin for ImageJ, using the timelapses described above. Only cells that did not move appreciably during the course of the movie were selected, as well as vesicles that could be clearly tracked throughout. Heatmap analysis was then performed using a custom script in R. Code is available at https://github.com/jkinglab/kinglabcode/blob/1cd29d61349f8a7cfd92274d31a8c795ecf6e47d/vesicle%20track%20heatmaps.R

### Statistical analysis

All data except heatmaps were analysed and plotted using Graphpad Prism 9. Each experiment consisted of at least 3 independent repeats. Where appropriate, biological replicates are plotted as dark symbols, with technical replicates (i.e. individual cells or vesicles) as faded symbols behind. Statistical analysis was performed on the mean values from each day, using an unpaired Mann-Whitney T-Test unless otherwise specified. Throughout * = P<0.01, **= P<0.005 and *** P<0.001. Error bars indicate standard error of the mean, unless otherwise specified.

## Acknowledgements

The authors are extremely grateful to Richard Kessin (Emeritus Professor, Colombia University Irving Medical Center, RichardKessin.com/) for sharing the unpublished hits of their elegant autophagy screen that started this project off. We also thank Pauline Schaap (University of Dundee) for sharing the *D. lacteum* genome and cells with us, as well as Ralf Graf (University of Potsdam) for both useful microtubule reagents and expert advice.

## Supplementary movie legends

**Supplementary Movie 1:** Autophagosome formation in wild-type and *katA*^*-*^*1* cells. Cells expressing GFP-Atg8aimaged by spinning disc after compression under agarose.

**Supplementary Movie 2:** Photo-induced relocation of GFP-dlKatnip. Confocal movie of wild-type cells under continuous illumination on a spinning disc microscope.

**Supplementary Movie 3:** Photo-induced relocation of GFP-dlKatnip after 30 µM nocodazole treatment. Cells were images as in movie 2. Note the more dense localisation to the shortened microtubules.

**Supplementary Movie 4:** Dynamics of GFP-α-tubulin in wild-type and *KatA*^*-1*^ cells. Maximum intensity projection from a confocal timelapse.

**Supplementary Movie 5:** Mitosis of wild-type cells, expressing GFP-α-tubulin.

**Supplementary Movie 6:** Mitosis of Katnip null cells expressing GFP-α-tubulin.

**Supplementary Movie 7:** Example of phagosome movement within control and *KatA*^*-1*^ cells. Cells were allowed to engulf 1µm beads for 10 minutes prior to compression under agarose and imaging by DIC microscopy.

---

**Supplementary Figure 1:**
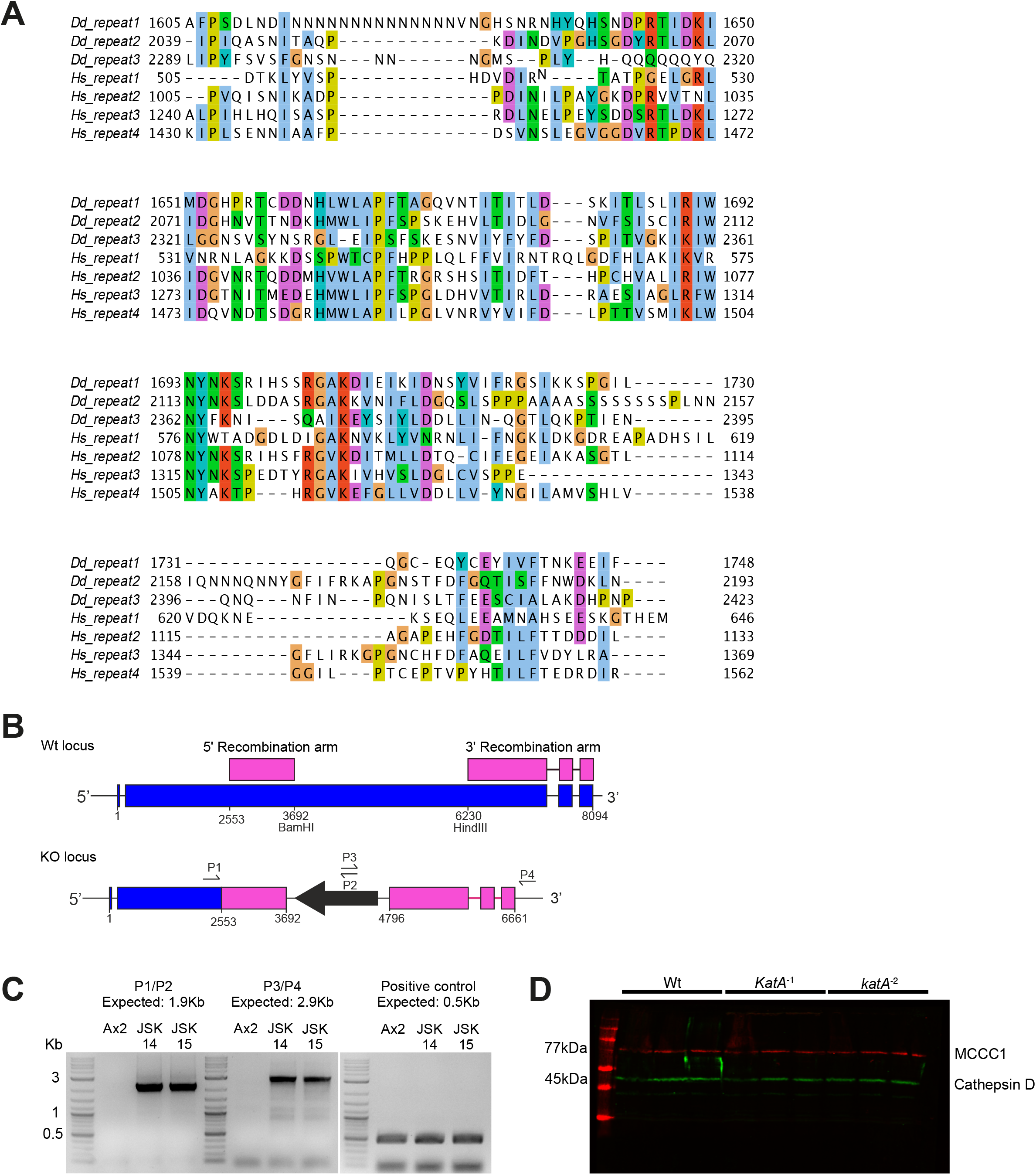
(A) CLUSTAL alignment of each DUF4457 repeat from the *Dictyostelium* and human Katnip orthologues. (B) Schematic of the knockout strategy showing the genomic locus before (top) and after *katA* disruption. P1-4 indicate the positions of primers used for screening for successful disruption by PCR. (C) Shows the PCR validation. Bands in the first two PCR’s indicate correct 5’ and 3’ arm intregration. The third PCR is a positive control. (D) Western blot of whole cell lysates from wild-type and *katA* mutant cells, probed for cathepsin D (green) and MCCC1 as a loading control. Three independent samples were run on the same gel for the quantification in Figure 3B.

**Supplementary Figure 2:**
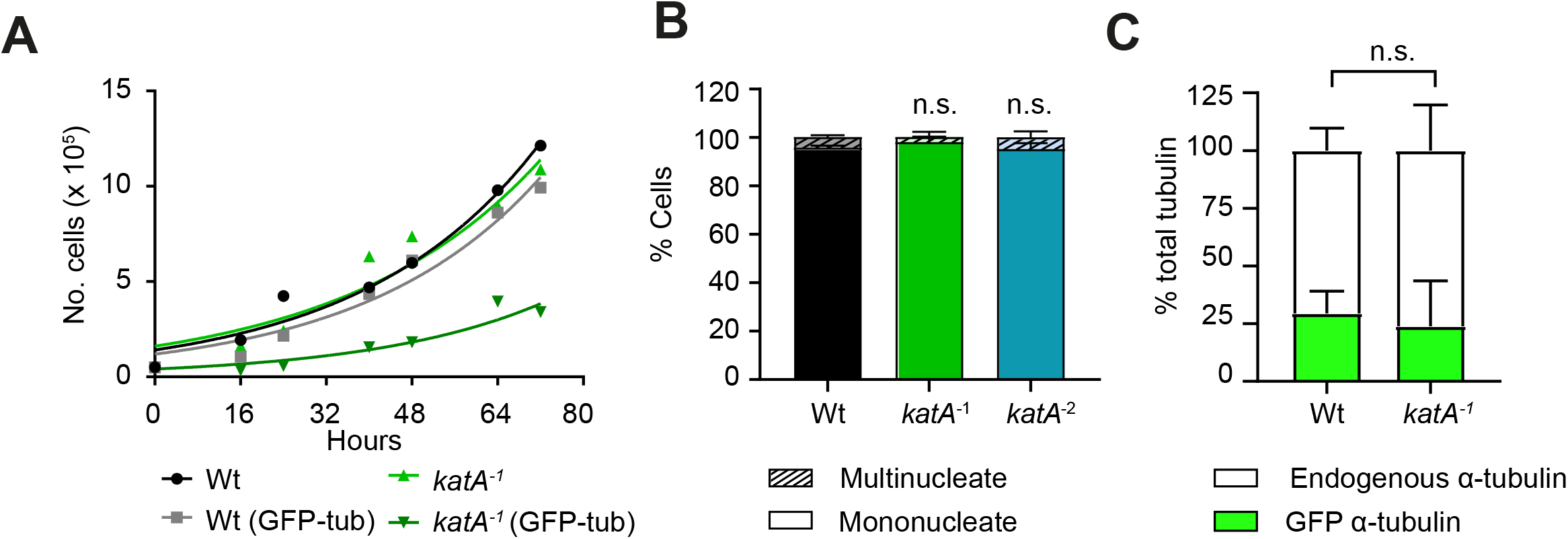
(A) Full growth curves of wild-type and *katA*^*-1*^ cells with and without expression of GFP-α-tubulin expression. (B) Quantification of the proportion of multinucleate cells. Cells were fixed and nuclei stained with DAPI. (C) Comparable GFP-α-tubulin expression levels in wild-type and *katA*^*-1*^ cells. Lysates were analysed by an α-tubulin western blot as in Figure 7F.

